# Unintentional seed dispersal via container-grown garden plants: Seed density and composition vary among exporter countries

**DOI:** 10.1101/2025.09.11.675287

**Authors:** Judit Sonkoly, Viktória Törő-Szijgyártó, Edit Petró, Attila Takács, Luis Roberto Guallichico Suntaxi, Andrea McIntosh-Buday, Patricia Elizabeth Díaz Cando, Katalin Tóth, Szilvia Madar, Gergely Kovacsics-Vári, Péter Török

**Affiliations:** Department of Ecology, University of Debrecen, Debrecen, Hungary; HUN-REN-UD Functional and Restoration Ecology Research Group, Debrecen, Hungary; HUN-REN-UD Conservation Biology Research Group, Debrecen, Hungary, Debrecen, Hungary; Polish Academy of Sciences, Botanical Garden - Centre for Biological Diversity Conservation in Powsin, Warszawa, Poland

**Keywords:** alien plants, container plants, contaminant pathway, horticultural trade, invasive species, plant nursery, plant trade

## Abstract

The global ornamental plant trade is well-known as an important source of alien plants, but the unintentional dispersal of seeds as contaminants of this trade is rarely considered and largely understudied. We sampled the substrate of imported container-grown plants in several garden centres in Hungary to answer the following questions: (i) What species’ seeds are present in the substrate of imported garden plants and in what quantity? (ii) Do the substrates of plants imported from different countries contain seeds in different density and composition? (iii) Do characteristics of the host garden plants affect the number and diversity of seeds found in their substrate? We detected altogether 2,181 seeds of 80 taxa in the substrates. On average, 1 litre of substrate contained 36 seeds of five species. Most species were alien in Hungary and considered invasive in Europe. The majority of samples contained at least one seed of a species invasive in Europe. We found that the substrate of plants imported from different countries contained seeds in different number, diversity, and composition. Thus, the structure of the horticultural trade network can shape the alien flora of importing countries. Furthermore, the substrate of needle-leaved ornamental species contained more seeds than the substrate of broad-leaved ornamentals. Our findings demonstrate that the ornamental plant trade can disperse a large number of seeds of numerous species and introduce alien species into the importing countries. We conclude that this introduction pathway deserves greater attention in invasion biology and more studies of this phenomenon are urgently needed.

## Introduction

Invasive species are among the most important factors impacting ecosystems worldwide (Díaz et al. 2019), having numerous negative effects on biodiversity and ecosystem functions (Pyšek et al. 2020), and even on human health and economy (Early et al. 2016, Haubrock et al. 2021). Global changes, especially climate warming will likely exacerbate the already vast problem posed by invasive species (Walther et al. 2009, Early et al. 2016); therefore, being able to understand and foresee biological invasions is more and more important in the era of global change (Bradley et al. 2010, Pyšek et al. 2020). The activities of humans are known to be most important factors driving biological invasions (Pyšek, et al. 2010, Bertelsmeier et al. 2017). Trade and transport shape invasions especially strongly (Perrings et al. 2005, Meyerson and Mooney 2007), with most harmful invasive species reaching new regions through trading activities (Levine and D’Antonio 2003). The monetary value of imports into a country is also clearly linked to the number of alien species in that country (Vilà and Pujadas 2001, Westphal et al. 2008). Therefore, the characteristics of trade and transport networks can even be used to assess invasion risk and the propagule pressure associated with specific commodities (Chapman et al. 2017). The volume of international trade is expected to increase in the coming decades (Chateau et al. 2015), likely resulting in a further increase in the number of alien species worldwide.

The likelihood of a species becoming invasive in a region is known to depend on the introduction pathway by which it reached the particular region (Hulme et al. 2008a, Pyšek et al. 2011). Information on what species are introduced via what pathways would therefore facilitate risk assessment and import regulations (Hulme et al. 2008b, Saul et al. 2017) and reduce alien species introductions (Hulme et al. 2008a, Novoa et al. 2020). The transport and introduction phases of invasions are clearly highly relevant for invasive species management and policy, yet these phases are less researched than the spreading phase and the effects of invasive species (Chapman et al. 2017). It was also emphasised by many authors that more effort and resources should be concentrated on the introduction stage of invasions (Leung et al. 2002, Westphal et al. 2008, Simberloff et al. 2013). Based on the above considerations, having a comprehensive view of the different pathways by which species are introduced into new regions would be vital (Essl et al. 2015, Chapman et al. 2017).

A high proportion of the alien plant species worldwide have been deliberately introduced as ornamental plants (Haeuser et al. 2018, Dawes et al. 2025), but unintentional introductions are also known to be a prominent source of alien invasive species (Lambdon et al. 2008, Lehan et al. 2013). In the Czech Republic for example, 772 plant taxa were listed as introduced unintentionally (Pyšek et al. 2022). Unintentionally introduced alien species frequently reach new regions as contaminants in traded commodities (Hulme et al. 2008a). According to Pyšek et al. (2009), 403 plant species have been introduced to Europe as contaminants of commodities (e.g., via seeds and mineral materials), which constitute 17% of all recorded alien species. Intentional releases and escapes from cultivation are well-recognized introduction pathways; therefore, their rates are already declining (Hulme et al. 2008a). On the other hand, the number of new introductions through other pathways (such as the contaminant, stowaway, and corridor pathways) is still increasing (Hulme et al. 2008a, Seebens et al. 2017).

Most studies demonstrating that invasions are closely linked to international trade considered trade as a whole, such as the total value of imports (e.g., Vilà and Pujadas 2001, Westphal et al. 2008). However, more recently, the live plant trade has been found to be more closely related to invasions than trade in general (Chapman et al. 2017). The international live plant trade experienced a significant growth in recent decades (Hinsley et al. 2025), and is a considerable part of global commerce, with an export value of USD 11.8 billion in 2024 (International Trade Center 2025). Horticultural trade, considering both live plants and horticultural substrates (growing media), has been shown to disperse a range of plant pathogens (e.g., Liebhold et al. 2012, Bienapfl and Balci 2014), invertebrates such as insects and snails (Sæthre et al. 2010, Moore et al. 2016), and even vertebrate species (Silva-Rocha et al. 2015, Rato et al. 2023). In certain areas, many different alien species have been first documented in plant nurseries and garden centres, for example snails (Bergey et al. 2014, Páll-Gergely et al. 2020) and planarians (Lazányi et al. 2024), which also suggests that they presumably reached the new sites via the horticultural trade. Several alien plant species have also been first detected in nurseries and garden centres (e.g., Hoste et al. 2009, Molnár V. et al. 2024, Sonkoly et al. 2024), but direct assessments of the dispersal of plant propagules as contaminants of horticultural stock are rare. The viable seed content of imported container-grown plants was studied in Alaska (Conn et al. 2008) and in Norway (Bruteig et al. 2017), and the unordered additional plant species and seeds accompanying aquatic plants ordered from plant vendors throughout the USA were assessed by Maki and Galatowitsch (2004). However, to our knowledge, there are no other studies of this issue so far.

Propagule pressure is considered one of the most important factors determining the success of biological invasions (Colautti et al. 2006, Lockwood et al. 2005); thus, deepening our understanding of it is vital for anticipating future invasions (D’Antonio et al. 2001, Cassey et al. 2004). Unfortunately, direct information on propagule pressure is difficult to obtain (D’Antonio et al. 2001, Eschtruth and Battles 2011), especially for plants and other organisms introduced via small propagules that are challenging to detect and identify. As the live plant trade is now recognised as a pathway for accidental alien plant introductions (mostly in the form of contaminant seeds), it is timely to increase our knowledge of the species dispersed via this pathway and the propagule pressure it generates. However, invasive species policy in ornamental horticulture generally overlooks the accidental dispersal of contaminant plant species (see e.g., Barbier et al. 2013, Hulme 2015, Hulme et al. 2018). Even a recent review summarising the environmental and societal risks of the international ornamental plant trade has ignored contaminant plant species and only focused on the animal species hitchhiking on traded live plants (Hinsley et al. 2025). Moreover, while deliberately importing soil is subject to strict regulations in many countries, these regulations do not apply to soil attached to plant roots or the substrate of imported container-grown plants, which may facilitate the large-scale spread of alien species (Hulme et al. 2008a).

Based on the above considerations, it is timely to understand what plant species are dispersed as contaminants of container-grown plants and in what quantities. Identifying the factors that influence seed density and composition of container substrates is also essential for improving our ability to anticipate introductions and to design more effective preventive measures and policy instruments. The exporter country of the plant may be such an influencing factor (see Bruteig et al. 2017). The substrates of container-grown plants arriving from different countries may contain different amounts of seeds and in different composition due to several reasons, including differences in their regional flora or in the weed management practices employed by the nurseries (Conn et al. 2008, Stewart et al. 2017). Some characteristics of the garden plants may also influence the number and diversity of seeds in the substrates (see e.g., Conn et al. 2008). For example, it usually takes more time to grow woody plants than herbaceous ones, which provides more time for seeds to accumulate in their substrates. Species with slow growth rates also take more time to reach a marketable size than fast-growing species, therefore they may also be more likely to accumulate seeds in their containers.

In this study, we sampled the substrate of recently imported container-grown garden plants in multiple garden centres in Hungary to answer the following questions: (i) What species’ seeds are introduced as contaminants of the substrate of garden plants and in what quantity? (ii) Do the substrates of garden plants imported from different countries contain seeds in different density and composition? (iii) Do characteristics of the garden plants affect the number and diversity of seeds found in their substrate?

## Materials & Methods

### Sampling and germination

To assess the viable seed content of the substrate of imported container-grown plants for garden use (hereafter ‘garden plants’), we took substrate (i.e., growing media) samples from the containers of recently imported perennial garden plants in six garden centres in North-Eastern Hungary in the spring of 2023 (Table 1, Fig. 1). We aimed to sample the substrate of recently imported plants to avoid collecting seeds and other propagules (hereafter referred to as ‘seeds’ for simplicity) that may have been accumulated in the containers due to exposure to the local seed rain. To this end, garden centre employees assisted us in selecting plant batches that had arrived recently. We collected a total of 60 one-litre samples (see Supplementary material Table S1). Each sample was pooled from multiple containers of the same garden plant species and batch, which were chosen based on the availability of batches that arrived recently. For large garden plants, whose containers were also large and available in limited numbers, the one-litre sample was collected from as few as two containers. In contrast, for small garden plants, samples were pooled from 10 to 20 containers, which also helped to avoid damaging the roots. In cases when the surface of the substrate was covered with a mulch layer (wood shavings or pine bark), we excluded the mulch layer from the sample. Employees also provided information about the country of origin of the plants: 42 of the sampled garden plant batches were imported from Italy, 16 from the Netherlands and two from Spain (see Supplementary material Table S1).

**Table 1.**
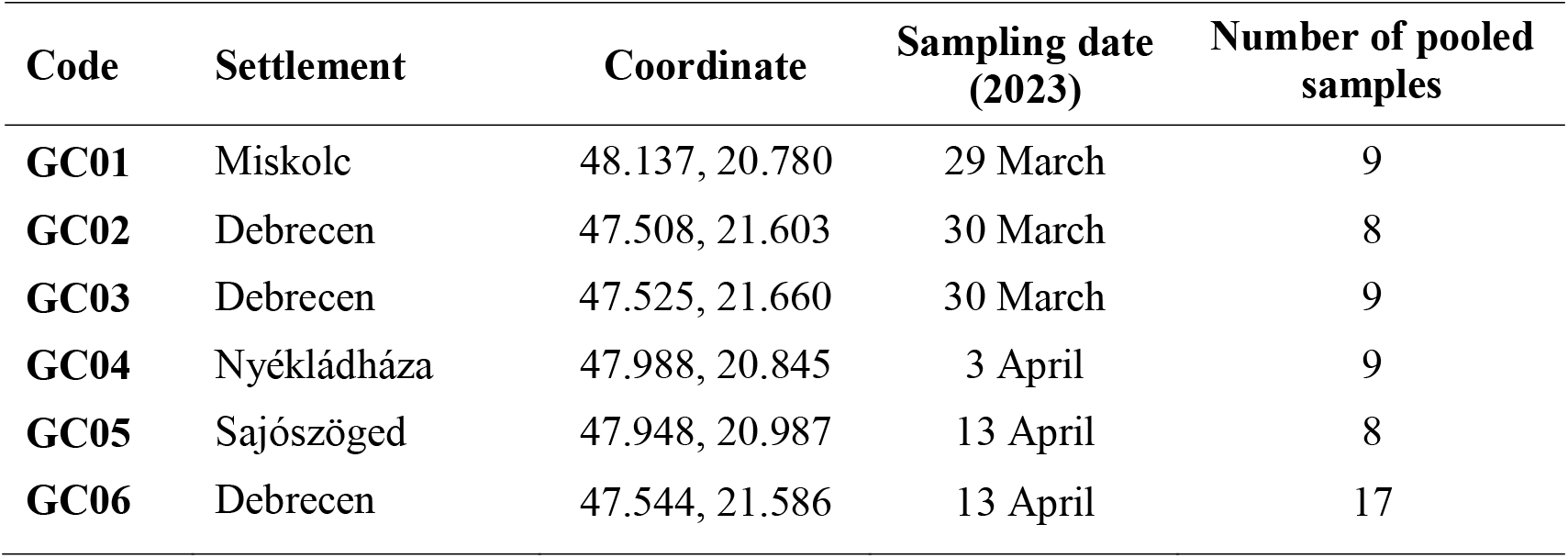
The location of sampled garden centres in NE Hungary, and the number of substrate samples taken from them. One pooled sample is 1 litre of substrate.

**Figure 1.**
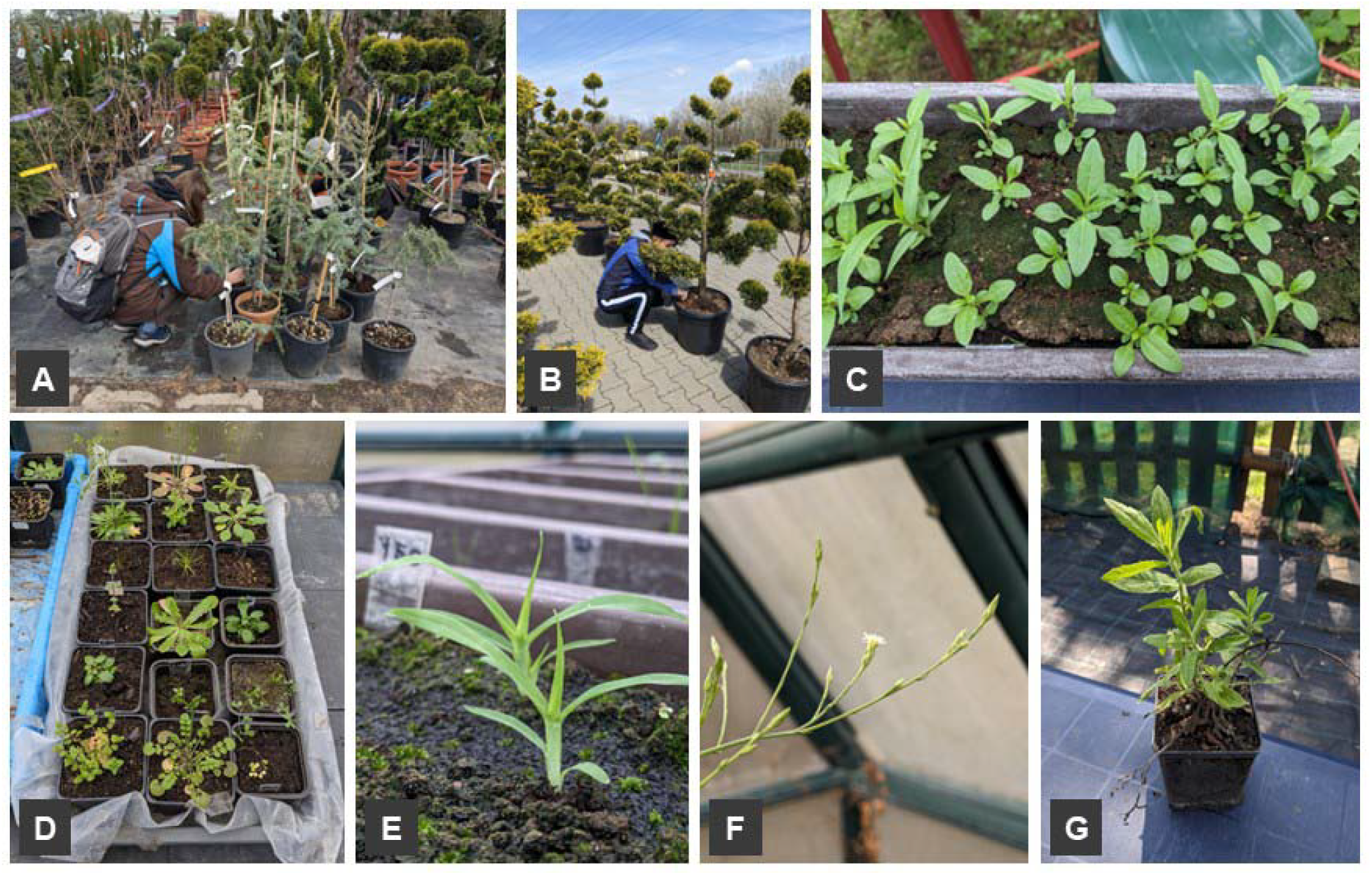
A– Sampling the substrate of a batch of *Cedrus deodara* ‘Feelin Blue’. **B** – Sampling the substrate of a batch of *Cupressocyparis leylandii* ‘Gold Rider’. **C** – *Eclipta prostrata* and a few other seedlings emerging from a substrate sample. **D** – Plants previously transplanted and waiting for identification. **E** – *Eleusine indica* emerging from a sample. **F** – *Symphyotrichum squamatum* germinated from a sample and flowering in the greenhouse. **G** – Individual transplanted for later identification (*Dittrichia viscosa*). All people depicted in the photographs are authors of the study who gave their consent for inclusion. Photographs by Judit Sonkoly.

The collected samples were then transported to the laboratory of the Department of Ecology, University of Debrecen (Hungary). We used the method of Ter Heerdt et al. (1996) to reduce sample volume, which involves washing the samples through two sieves (mesh size 0.2 mm and 2.8 mm). This method eliminates fine particles by washing them out, while large particles are retained by the coarse mesh, considerably reducing sample volume. The concentrated samples were then spread on the surface of steam-sterilised potting soil in balcony planters (sized approximately 60 cm × 20 cm × 15 cm). Each balcony planter contained a one-litre sample and we also set up five planters only containing sterilised potting soil (no sample) to detect seed contamination during the germination period. Therefore, altogether 65 balcony planters were used in the experiment, which were kept in an unheated greenhouse in the botanical garden of the University of Debrecen. The germination period lasted from 17 April 2023 to 16 November 2023, during which the samples were watered on a daily basis. Emerging seedlings were regularly counted, identified and removed. We transplanted the plants which were not yet identifiable into separate pots. The transplanted plants (155 individuals) were further grown until they could be identified, but not later than November 2024, i.e., the end of the second vegetation season.

### Calculations and analyses

The number of seeds and the number of species per sample were compared between samples originating from Italy and the Netherlands using Wilcoxon rank sum tests. As only two samples were from garden plants imported from Spain, those two samples were excluded from analyses based on the country of origin, and only samples originating from Italy and the Netherlands were compared. Samples collected in different garden centres were compared regarding seed numbers and species numbers using Kruskal-Wallis tests and Dunn tests with Bonferroni correction as post hoc tests.

To assess differences in seed community compositions between samples originating in different countries and between samples collected in different garden centres, we performed permutational multivariate analyses of variance (PERMANOVAs) with Bray-Curtis dissimilarity, using the *adonis2* function from the *vegan* package in R. We assessed homogeneity of multivariate dispersions using the *betadisper* function in the *vegan* package, followed by a permutation test (999 permutations). No significant differences in dispersion were found, validating the PERMANOVA results. Principal coordinates analysis (PCoA) was used to visualise the compositional differences between samples originating in different countries.

Indicator species analysis was conducted using the *multipatt* function from the *indicspecies* R package (Cáceres and Legendre, 2009), applying the group-equalized IndVal.g method (Dufrêne and Legendre, 1997) with 999 permutations. To reduce noise from rare species in the sparse dataset, species occurring in less than three samples were filtered out before the indicator species analyses. After this filtering, 33 species were included in the indicator species analysis regarding countries, and 35 species were included in the analysis regarding garden centres.

To assess the effect of the garden plants’ taxonomy on the number and diversity of seeds found in their substrates, we categorized the garden plants into families based on the Plants of the World Online database (POWO 2025, see Supplementary material Table S1). Samples of the substrates of plants belonging to families represented by less than five species in our dataset were excluded from this analysis. We then compared seed numbers and species numbers in the substrates of garden plants belonging to different families using Kruskal-Wallis rank sum tests. As the garden plants were clearly separable into two large groups based on leaf habit: (i) needle or scale-leaved species (Conifers), and (ii) broadleaved species (Angiosperms, mostly dicots), we also tested whether the leaf habit of the garden plant had an effect on the number and diversity of seeds in its substrate using Wilcoxon rank sum tests.

All analyses were carried out in R version 4.3.2 (R Core Team, 2023). Nomenclature follows Euro+Med PlantBase (http://www.europlusmed.org). The native versus alien status of species and their invasion status in Hungary was given according to the Pannonian Database of Plant Traits (PADAPT, Sonkoly et al. 2023). Invasiveness in Europe is based on Kalusová et al. (2024); we considered a species as invasive in Europe if it is invasive in at least one region of Europe.

## Results

In total, the samples contained 2,181 germinating seeds. Fourteen seedlings died before they could have been identified (<1% in total, one monocot and 13 dicots), the further 2,167 seedlings belonged to 80 taxa, 74 of them identified at the species level (Supplementary material Table S2). Most of the individuals were identified on the species level, except for one *Cerastium*, three *Digitaria*, five *Juncus*, two *Trifolium*, and 8 *Typha* seedlings which died before further identification was possible or did not develop diagnostic features by the end of the second vegetation season (November 2024). *Juncus ranarius* and *J. bufonius* seedlings were pooled as *Juncus bufonius/ranarius* because they died before they could have been distinguished without doubt. The number of seedlings from one-litre samples ranged between 0 and 783, with a mean of 36.35 seedlings/L. The number of taxa germinated from one-litre samples ranged between 0 and 22, with a mean of 5.38 taxa/L (excluding the unidentified individuals, see Supplementary material Tabe S1).

The most abundant species were *Sagina procumbens* (957 seedlings), *Cardamine occulta* (221 seedlings), *Euphorbia maculata* (201 seedlings), *Portulaca oleracea* (77 seedlings), and *Oxalis corniculata* (76 seedlings) (Supplementary material Table S3). On the contrary, the majority of taxa were represented by 1–5 individuals. The majority of taxa were present in 1–5 samples (61 species), while the most common ones were *E. maculata* (in 31 samples), *P. oleracea* (in 19 samples), and *S. procumbens* (in 17 samples, see Supplementary material Tabe S3). There were seven species (*Digitaria sanguinalis, Erigeron canadensis, Euphorbia maculata, Oxalis corniculata, Poa annua, Portulaca oleracea*, and *Sonchus oleraceus*) which we detected in samples originating from all three countries, even though only two samples originated from Spain. Half of the taxa were detected in samples collected in a single garden centre, but five species were detected in samples from all garden centres (*C. occulta, E. maculata, P. oleracea, S. procumbens*, and *S. oleraceus*, see Supplementary material Tabe S3).

The majority of the detected species were alien in Hungary (48 species, Supplementary material Table S3). Six of the species we detected have only recently been documented in the country, while two species (*Dittrichia viscosa* and *Symphyotrichum squamatum*) have not yet been reported from Hungary. On average, 78.8% of the seeds detected in each sample belonged to species alien in Hungary, and 91.7% of the samples contained at least one seed of an alien species. On average, 28.6% of the seeds detected in each sample belonged to species considered to be invasive in Hungary, and 73.3% of the samples contained at least one seed of a species invasive in Hungary. Regarding their invasiveness in Europe, 53.9% of the seeds detected belonged to species considered to be invasive in at least one region of Europe and 86.7% of the samples contained at least one seed of a species invasive in Europe.

Both the number of seeds and the number of species were significantly higher in samples originating from Italy compared to samples originating from the Netherlands (W=160, *p*=0.002, and W=154, *p*=0.001, respectively, Fig. 2). The garden centre where the sample was collected also had a significant effect on both seed number per sample (χ^2^(5)=17.422, *p*=0.004) and species number per sample (χ^2^(5)= 15.141, *p*=0.010). However, pairwise post hoc tests indicated a significant difference in only two of the 15 garden centre pairs for both seed number and species number. Seed and species numbers per sample at the remaining garden centres did not differ significantly from each other (Supplementary material Table S4 and S5). PERMANOVA revealed that both the country of origin (F=2.541, R^2^=0.046, *p*=0.001, Fig. 3) and the garden centre where the sample was collected (F=1.431, R^2^=0.123, *p*=0.004) had a significant effect on the seed composition of the samples.

**Figure 2.**
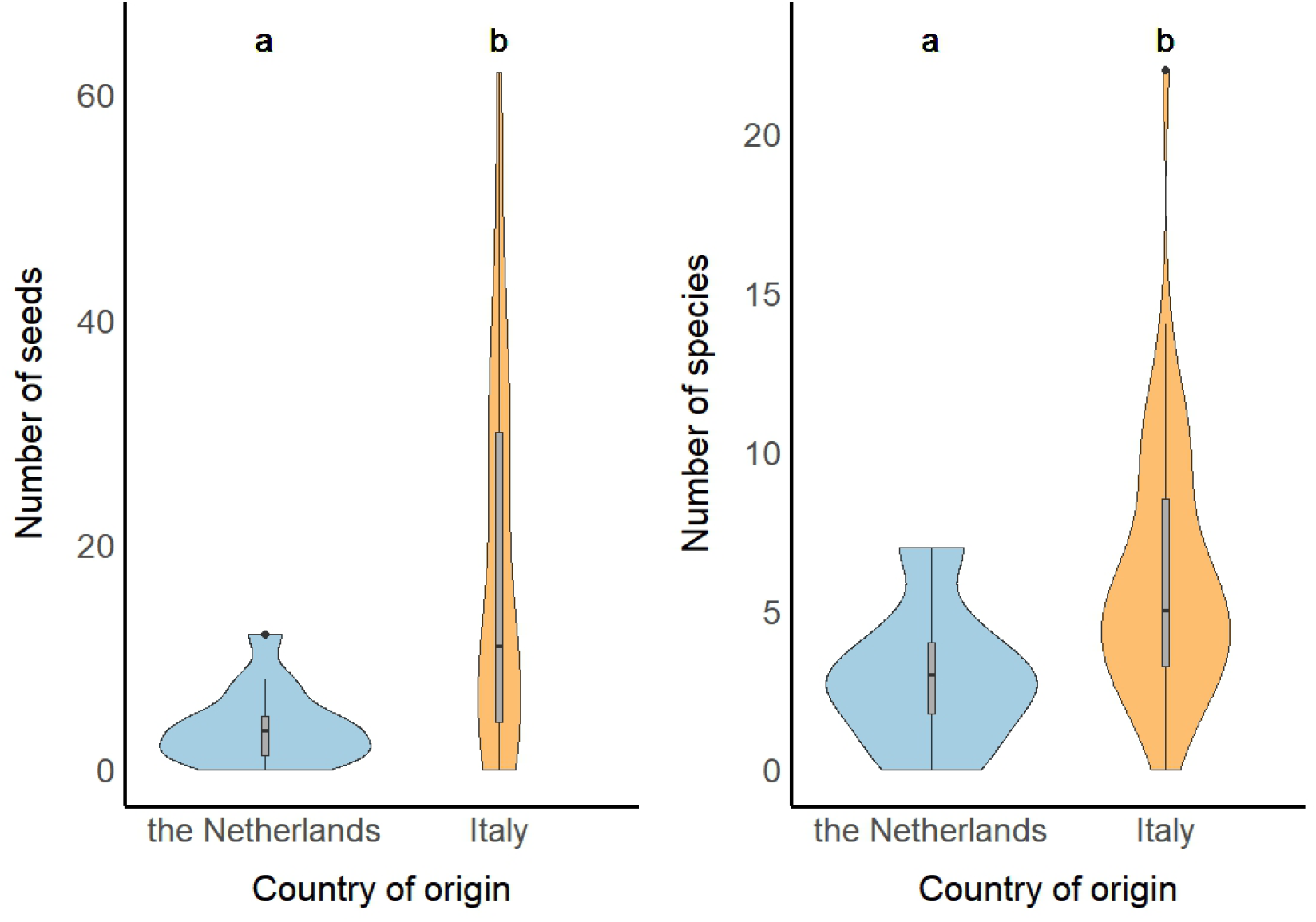
The number of seeds and the number of species in one-litre samples taken from the substrate of garden plants imported from the Netherlands (n=16) and from Italy (n=42). Note that outlier values are not shown on the seed number plot to improve the readability of the plot.

**Figure 3.**
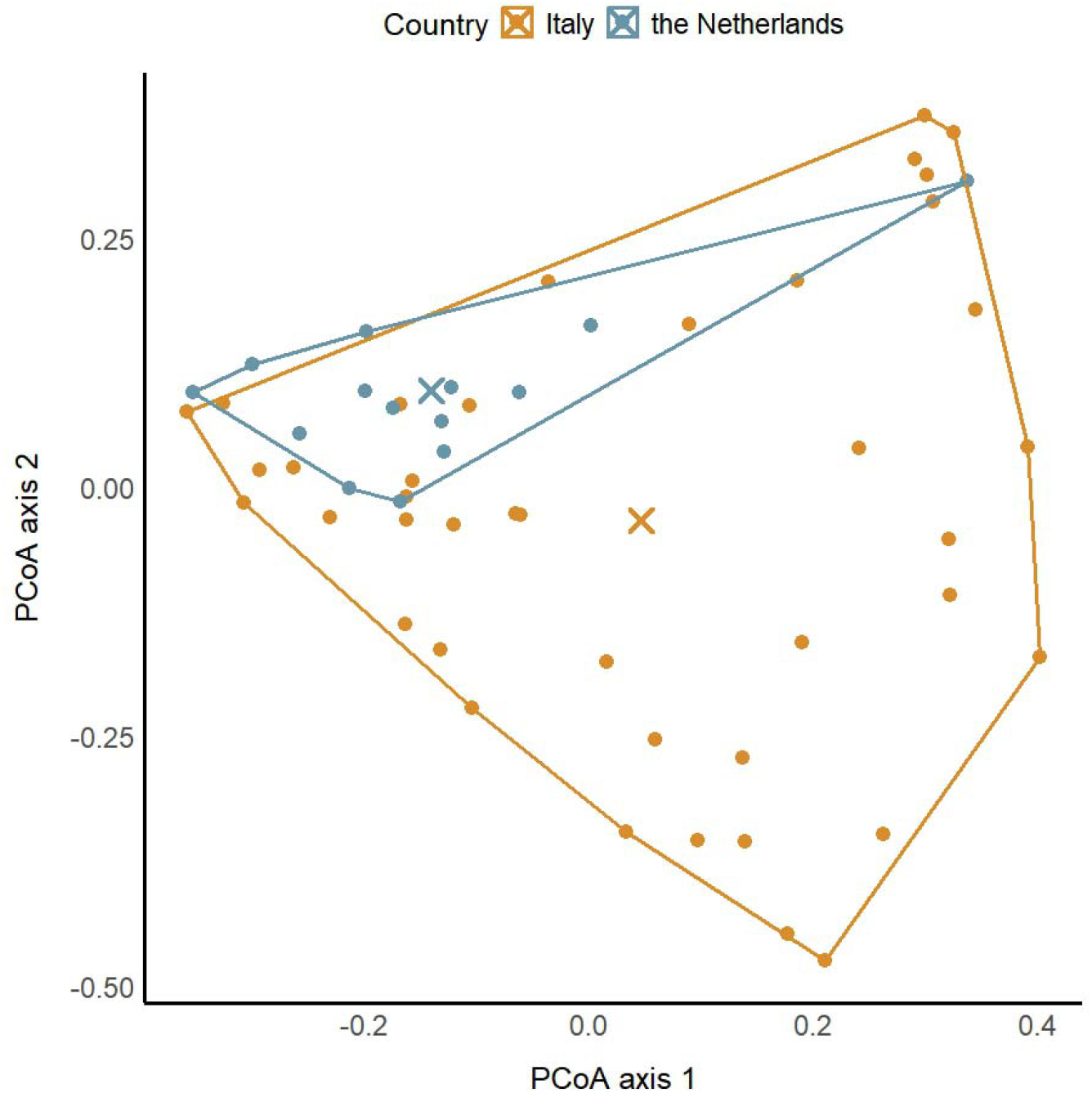
Principal Coordinates Analysis (PCoA) of the seed composition of samples by country of origin. The first two PCoA axes explained 12.1% and 9.5% of the variation in community composition, respectively. The × symbols indicate the centroids of the two groups.

Indicator species analysis identified four taxa significantly associated with one of the countries. *Euphorbia maculata* (IndVal=0.835, *p*=0.001), *Portulaca oleracea* (IndVal=0.631, *p*=0.034), and *Euphorbia prostrata* (IndVal=0.541, *p*=0.045) were associated with ornamental plants originating from Italy. *Cerastium glomeratum* was associated with ornamentals originating from the Netherlands (IndVal=0.499, *p*=0.019). The indicator species analysis for garden centres identified four taxa significantly associated with one or more of the garden centres where the samples were collected. *Chenopodium album* was an indicator of samples from the GC03 site (IndVal=0.62, *p*=0.011), *Amaranthus emarginatus* with the GC01 site (IndVal=0.62, *p*=0.01), *Sagina procumbens* (IndVal=0.78, *p*=0.011) with the group GC02 *+* GC04, while *Euphorbia maculata* showed a strong association with samples from the combined group GC02 *+* GC04 *+* GC06 sites (IndVal=0.808, *p*=0.009).

The family of the host garden plant did not have a significant effect on either the number of seeds (χ^2^(3)= 7.284, *p*=0.063) nor the number of species (χ^2^(3)= 5.094, *p*=0.165) detected in the samples (Supplementary Figure S1). The leaf habit of the host garden plant had a significant effect on the number of seeds (W=275, *p*=0.013), but not on the number of species detected in the samples (W=317.5, *p*=0.063, Fig. 4).

**Figure 4.**
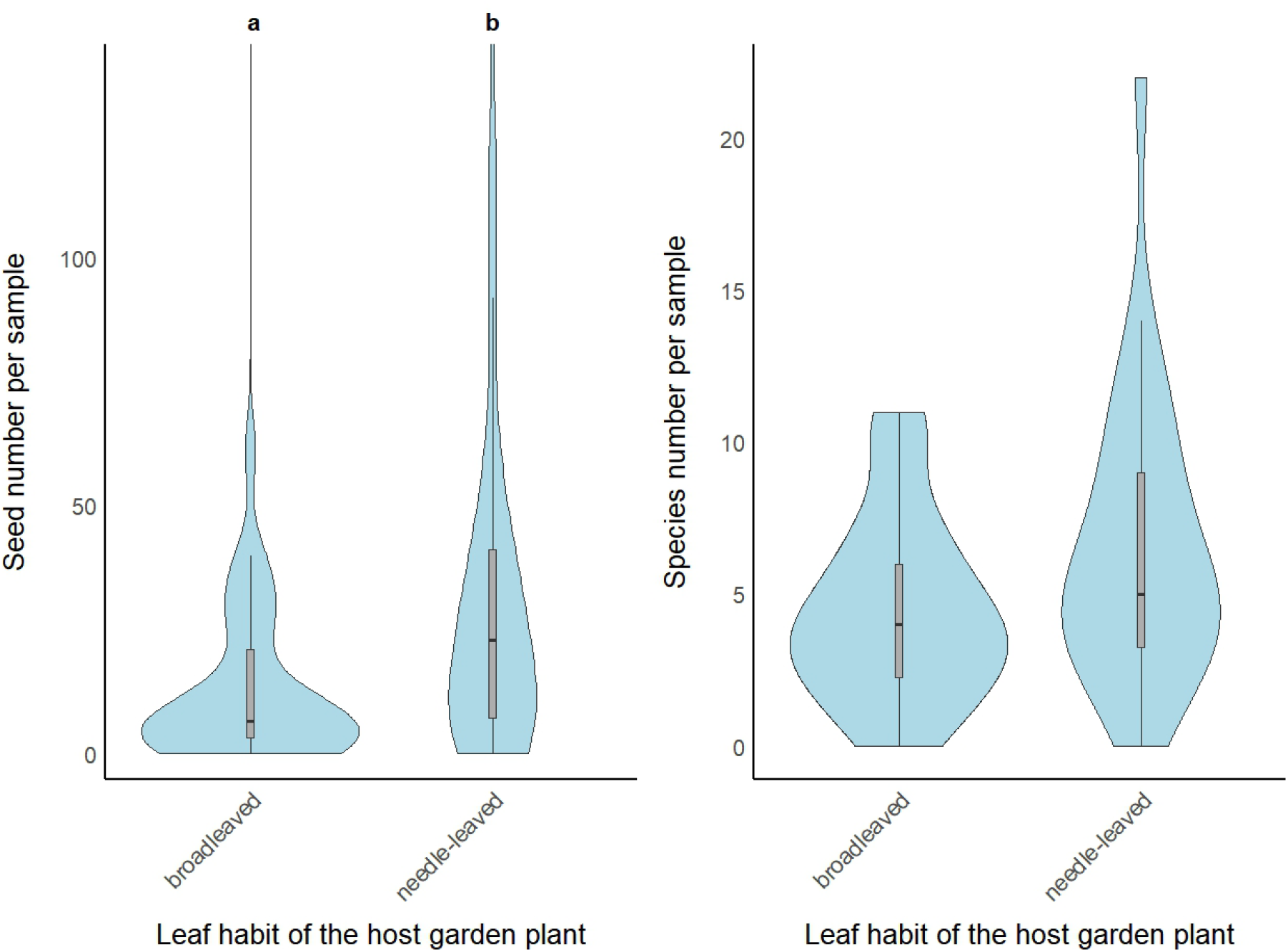
The number of seeds and the number of species in one-litre samples taken from the substrate of broadleaved (n=34) and needle-leaved garden plants (n=26). Note that outlier values are not shown on the seed number plot to improve the readability of the plot.

## Discussion

Human activities such as trade and transport have long been recognised as the main driving forces of alien species introductions and biological invasions (Hulme 2009, Bullock et al. 2018). Ornamental plants are known to be the main sources of naturalised and invasive plant species worldwide (Pergl et al. 2016, van Kleunen et al. 2018). In contrast, the accidental dispersal of contaminant seeds associated with the international live plant trade has not been given considerable attention so far. Here we showed that the substrate of container-grown garden plants imported from distant countries can contain large numbers of seeds – mostly of alien plant species. Moreover, the seed density and composition of the substrates can differ among exporter countries.

Nurseries producing container-grown ornamental plants are exposed to intensive disturbance and high propagule pressure due to the constant inflow of potting substrates and other horticultural materials. Resource availability is also high in these man-made habitats due to fertilizer use. All these factors can contribute to the high invasibility of these habitats (Davis et al. 2000, Davis and Pelsor 2001, Colautti et al. 2006); therefore, it is not surprising that they have been repeatedly found to be inhabited by numerous alien species (see e.g., Conn et al. 2008, Hoste et al. 2009, Sonkoly et al. 2024). Although most of the alien species germinated from the substrates are already widespread throughout Europe (e.g., *Erigeron canadensis, Galinsoga quadriradiata, Oxalis corniculata*), some of the species we detected are still in the early stages of invasion in most of the continent. For example, *Gnaphalium pensylvanicum* is a species native to America, whose spread has only been documented in Western and Southwestern Europe so far (Verloove et al. 2023). The only documented occurrence of *G. pensylvanicum* in Hungary is from the container of an ornamental plant in a garden centre (Sonkoly et al. 2024). *Dittrichia viscosa* is native to the Mediterranean Basin and has been introduced to some other parts of Europe, but it is not yet considered as an invasive species in most European countries (Kalusová et al. 2024) and it has not yet been reported from Hungary. *Symphyotrichum squamatum*, a species native to South America, is already considered invasive in several European countries (see e.g., Arianoutsou et al. 2010, Uludağ et al. 2017), but it mainly spreads in Southern Europe and has not been reported from Hungary yet. Some seeds of ornamental plant species were detected as well, but in surprisingly low numbers: three seedlings of *Nerium oleander* germinated from the substrate of the same species and one seedling of *Lonicera nitida* was detected in the substrate of *Taxus* × *media* ‘Hillii’ individuals.

There were seven species (*Digitaria sanguinalis, Erigeron canadensis, Euphorbia maculata, Oxalis corniculata, Poa annua, Portulaca oleracea, Sonchus oleraceus*) which we detected in samples originating from all three countries, even though there were only two samples from Spain. These species frequently grow in the containers of ornamental plants according to our experiences during sample collection, as well as to previous results (Sonkoly et al. 2024). 45 of the species we documented (60.8%) were also detected in samples from the containers of plants imported to Norway (Bruteig et al. 2017). The plant species which were represented by the highest numbers of seeds in our study also considerably overlap with the species Bruteig et al. (2017) reported as the most abundant species germinating from the samples. Case et al. (2005) listed 26 species as common nursery weeds in the USA, 14 of which were also found in our study. The detected alien species also show a great overlap with the alien species detected in a survey assessing the alien flora of nurseries and garden centres in Hungary (Sonkoly et al. 2024). Based on these findings, we can presume that there is a limited set of plant species which are particularly successful in colonizing plant nurseries in Europe and the USA, and therefore able to build up a considerable seed bank inside the containers.

The average number of seeds per litre of substrate in our study (36.4 seeds/L) was considerably higher compared to the seed densities found by Conn et al. (2008) in Alaska. They found the highest number of seeds in the soil of balled-and-burlapped trees and shrubs (20.2 seeds/L), followed by small and large container-grown woody plants (7.8 seeds/L and 3.2 seeds/L, respectively). The number of taxa detected per litre was also much higher in our study. Conn et al. (2008) found the highest number of species in the soil of large container-grown woody plants (1.8 species/L), while we found an average of 5.4 species/L. A report by Bruteig et al. (2017) assessed the seed content of the substrate of container plants imported to Norway. They reported a total of 17,123 seedlings from 650 litres of samples (26.3 seeds/L), which is comparable to the seed density in our study, although they did not provide the number of seeds per litre directly.

The substrates of plants imported from Italy contained seeds in significantly higher density and diversity than the substrates of plants imported from the Netherlands. Substrates of plants imported from different countries also had significantly different seed compositions, although the variance explained was rather low. Species documented exclusively in the substrates of plants imported from the Netherlands were almost all native and widely distributed throughout Europe (e.g., *Hypochaeris radicata, Juncus inflexus, Urtica urens*). Conversely, many species found only in the substrates of Italian plants are primarily distributed in Southern Europe. Some of these are native to the region (*Dittrichia viscosa, Polypogon virdis*), while others are alien species that have mainly colonised Southern Europe so far (e.g., *Eclipta prostrata, Euphorbia prostrata*, and *Symphyotrichum squamatum*). Although further studies are definitely needed to corroborate our findings, they suggest that the substrate of garden plants imported from different countries can be considerably different in terms of seed numbers and species composition, which was also observed by Bruteig et al. (2017). Different countries have different floras and environmental conditions, and the weed management practices applied by nurseries may also vary between countries (Conn et al. 2008, Stewart et al. 2017), likely contributing to the observed differences in seed contamination between countries. These differences may also be related to the different range of species imported from the two countries. Plants imported from Italy mostly belonged to the Cupressaceae and Pinaceae families, while plants imported from the Netherlands belonged to a more diverse range of families, but were typically not Conifers (see Supplementary Table S1). Furthermore, substrate mixes are likely already contaminated with seeds before potting (see e.g., Case et al. 2005, Sonkoly et al. 2022), and different nurseries and countries may use different types of substrates.

Samples taken from different garden centres also had significantly different seed compositions. As the sampled garden plants were freshly imported, there was little opportunity for local seed contamination or for ecological filtering by the garden centres’ environment, suggesting that another external factor was likely responsible for the difference. It may have arisen from the fact that there were garden centres where we could obtain samples from garden plants imported from a single country, resulting in an association between garden centres and exporter countries. This is in line with our finding that only one garden centre (GC05) differed from some others in terms of both seed numbers and species numbers, and all the sampled garden plants were exported from the Netherlands in that garden centre.

Although the number and species diversity of seeds found in the substrates did not differ between garden plants from different families, the host garden plants’ leaf habit appeared to be associated with differences in seed content. It is well-established that species with needle or scale-like leaves have low specific leaf area (SLA; e.g., Reich et al. 1995, 1997), which is characteristic to species with a resource-conservative strategy and low growth rates (Shipley 2006, Poorter et al. 2009), while broadleaved species generally have higher SLA and grow faster. Therefore, the leaf habit of the garden plants may be associated with the length of time they spend in the nursery before being sold at a retail, during which time seeds can accumulate in their substrate (Conn et al. 2008), possibly explaining the observed differences.

The number of individuals introduced into a new area is typically quantified by propagule pressure, which is a measure considering both the number of individuals involved in separate introduction events (termed propagule size) and the number of such introduction events (propagule number) (Lockwood et al. 2005). Our results show that the propagule size component of propagule pressure can be rather high at least for some of the species dispersed in the substrates of traded garden plants. Species whose seeds are present in the substrates in high numbers (i.e., large propagule size) must have abundant seed sources within or at least nearby the source nurseries. Therefore, as container grown garden plants are regularly transported between the same nurseries and garden centres, we can presume that the propagule number component is also high for such species, resulting in high propagule pressure for many alien plant species. As garden centres are disturbed, resource-rich ‘habitats’ typically embedded in an intensively used, urbanized landscape and such habitats have been shown to have high invasibility (Davis et al. 2000, D’Antonio et al. 2001), if they are exposed to high propagule pressure of certain plant species, they are very likely to become starting points of invasions (Lockwood et al. 2005, Vedder et al. 2021).

The global horticultural trade has experienced considerable growth over the last two decades, with the export value of live plants approximately doubling during this time (Hinsley et al. 2025). In Norway, for example, the imports of container-grown garden plants have quadrupled between 1997 and 2016 (Bruteig et al. 2017). Italy and the Netherlands are the largest exporters of live plants globally (Hinsley et al. 2025). In 2021, the Netherlands exported almost 300 million kilograms of garden plants, mostly to other European countries (de Waart et al. 2025). Given that the substrates of these plants likely make up a considerably proportion of this mass, a vast number of seeds are undoubtedly dispersed accidentally with them each year. Due to the huge amount of incoming seeds and the diverse array of species, it is highly likely that some species introduced with imported garden plants will become naturalised in the new area (see also Hoste et al. 2009).

Climate change is expected to have little effect on the establishment rate of new alien plants because of the stronger effect of propagule pressure (Hulme 2017). However, already introduced alien species previously limited by unfavourable climatic conditions might be able to increase their abundance and become invasive under changing climate (Chapman et al. 2014, Hou et al. 2014). Some species introduced via container-grown plants imported from the Mediterranean region are probably not able to form self-sustaining populations in Central Europe at present due to climatic limitations (Beaury et al. 2023). They may not be able to reproduce at all due to the climatic limitations, or they may only be viable under special microclimatic conditions (Géron et al. 2021). However, further climate warming is likely to reduce existing climatic limitations for such species, facilitating their naturalisation and potential invasion (Walther et al. 2009, Dullinger et al. 2017). Therefore, the large number and diversity of seeds entering Central Europe as contaminants of container-grown plants from Italy (and possibly other Mediterranean countries) will be even more likely to trigger invasions as temperature rises (Seebens et al. 2015).

Analyses show that policy measures aimed at controlling invasions can effectively reduce the establishment of new alien species (Canelles et al. 2025). However, to design such measures, information on the risks posed by the unintentional introduction of species as contaminants of particular commodities is essential, as it can inform policy and help limit future invasions (Essl et al. 2015). Consequently, there is a pressing need for studies that assess which species are introduced via specific commodities and the associated propagule pressure. Increasing evidence shows that biological invasions are largely driven by global trade networks (e.g. Levine & and D’Antonio 2003; Westphal et al. 2008; Chapman et al. 2017), and detailed information on these networks is expected to improve risk assessments and support the development of effective policies (Banks et al. 2015; Paini et al. 2016). Improving our knowledge of which commodities are responsible for transporting specific contaminant species could therefore enhance the accuracy of invasion predictions. In particular, the live plant trade has been shown to be more strongly associated with invasions than trade in general (Chapman et al. 2017). Data on species accidentally transported via horticultural trade could facilitate more precise risk assessments (Essl et al. 2015), as the trade volume of horticultural commodities may serve as a proxy for propagule pressure for these species (Chapman et al. 2017). On the other hand, detailed and up-to-date information on the volume and spatial structure of the live plant and horticultural trade would also be needed to quantify propagule pressure (Colautti et al. 2006) and assess the risk posed by alien species dispersed via these commodities (Hulme 2009). These considerations highlight the need to pay greater attention to the accidental introduction of species as contaminants in the global horticultural trade.

In conclusion, the great number of seeds entering new regions as contaminants of the horticultural trade, most of which are alien species, is likely to induce new invasions due to high propagule pressure. We found that container-grown plants imported from different countries introduced seeds in different numbers and composition, suggesting that the structure of the horticultural trade network can shape the alien flora of importing countries and have important consequences for biological invasions. Characteristics of the imported container plants may also have an effect on the number of seeds dispersed in their substrate, but the effect of exporter country seems to be more important. Finally, our findings suggest that although the global horticultural trade has mostly been considered as a pathway for deliberate alien plant introduction so far, increased attention to the accidental dispersal of seeds as contaminants in horticultural trade is urgently needed. We also emphasize that more effort and research needs to be devoted to the less researched transport and introduction phases of biological invasions in general.

## Acknowledgements

We are grateful to the garden centres’ management for making sample collection possible, and to the employees for their help during sampling. We would also like to thank Tamás Wirth for confirming the identification of *Erigeron bonariensis*.

## Funding

The authors were supported by the National Research, Development and Innovation Office [J.S.: PD 137747; P.T.: KKP 144068, K 137573] during the manuscript preparation. V.T.-S. was supported by the University Research Scholarship Programme of the National Research, Development and Innovation Office (EKÖP-24-3-II-DE-220). The work of J.S. was also supported by the Bolyai János Scholarship of the Hungarian Academy of Sciences [BO/00587/23/8].

## Competing Interests

The authors have no relevant financial or non-financial interests to disclose.”

## Author contributions

J.S.: Conceptualization, Methodology, Investigation, Formal analysis, Visualization, Writing - Original draft, Funding Acquisition; V.T.-S.: Investigation, Resources, Project administration, Writing - Review and Editing; E.P.: Investigation, Writing - Review and Editing; A.T.: Investigation, Writing - Review and Editing; L.R.G.S.: Investigation, Writing - Review and Editing; A.M.-B.: Investigation, Writing - Review and Editing; P.E.D.C.: Investigation, Writing - Review and Editing; K.T.: Investigation, Writing - Review and Editing; S.M.: Investigation, Writing - Review and Editing; G.K.-V.: Investigation, Writing - Review and Editing; P.T.: Conceptualization, Writing - Review and Editing, Funding Acquisition.

## Supplementary Information

**Supplementary Table S1:** Information about the substrate samples (sampling date and location, ornamental plant, number of seeds germinated, number of taxa detected, etc.).

**Supplementary Table S2:** Detailed results of the germination study (species × sample matrix).

**Supplementary Table S3:** Detailed information about the detected species (number of seeds germinated, invasions status, etc.).

**Supplementary Table S4 and S5**: Results of the post hoc Dunn’s tests comparing seed numbers (Table S4) and species numbers (Table S5) in substrate samples collected in different garden centres.

**Supplementary Figure S1:** Violin plots comparing the number of seeds and the number of species in substrate samples taken from ornamental plant species belonging to different families.

## Notes

### Competing Interest Statement

The authors have declared no competing interest.

