## Supplemental figure S1 for "Unintentional seed dispersal via container-grown garden plants: Seed density and composition vary among exporter countries"

### Supplementary material

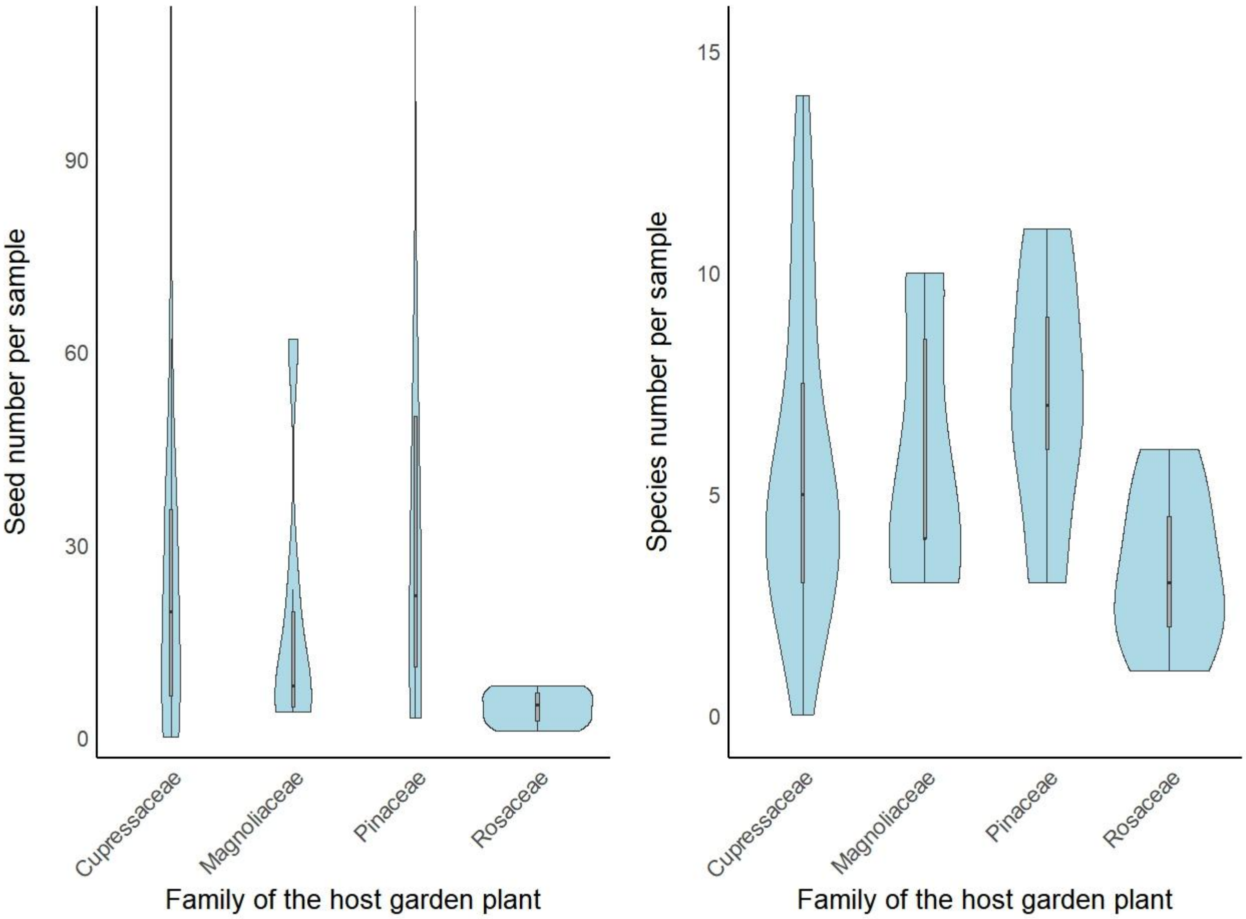


**Figure S1.** The number of seeds and the number of species in 1-litre samples taken from the substrate of species belonging to different families. Only families represented by at least five samples were compared. Note that outlier values are not shown on the seed number plot to improve the readability of the plot. Kruskal-Wallis rank sum tests revealed no significant differences in either case.
