## Supplemental tables S4 and S5 for "Unintentional seed dispersal via container-grown garden plants: Seed density and composition vary among exporter countries"

### Supplementary material

**Table S4.** Results of Dunn's test comparing seed numbers in samples collected from different garden centres (Z-score and Bonferroni-adjusted p-value). For details of the samples see Table 1.

| Sample code | GC01 | GC02 | GC03 | GC04 | GC05 |
|---|---|---|---|---|---|
| **GC02** | 0.533<br>1.000 | | | | |
| **GC03** | -2.216<br>0.400 | -2.683<br>0.1095 | | | |
| **GC04** | -0.797<br>1.000 | 1.306<br>1.000 | -1.419<br>1.000 | | |
| **GC05** | -3.059<br>*0.033** | -3.490<br>*0.007** | -0.909<br>1.000 | -2.285<br>0.334 | |
| **GC06** | -1.247<br>1.000 | 1.802<br>1.000 | -1.288<br>1.000 | 0.335<br>1.000 | -2.268<br>0.350 |

**Table S5.** Results of Dunn's test comparing species numbers in samples collected from different garden centres (Z-score and Bonferroni-adjusted p-value). For details of the samples see Table 1.

| Sample code | GC01 | GC02 | GC03 | GC04 | GC05 |
|---|---|---|---|---|---|
| **GC02** | 0.174<br>1.000 | | | | |
| **GC03** | -2.193<br>0.425 | -2.301<br>0.321 | | | |
| **GC04** | -1.106<br>1.000 | 1.248<br>1.000 | -1.086<br>1.000 | | |
| **GC05** | -3.019<br>*0.038** | -3.103<br>*0.029** | -0.892<br>1.000 | -1.946<br>0.775 | |
| **GC06** | -0.972<br>1.000 | 1.132<br>1.000 | -1.535<br>1.000 | -0.293<br>1.000 | -2.487<br>0.193 |
